# Identification of biological mechanisms underlying a multidimensional ASD phenotype using machine learning

**DOI:** 10.1101/470757

**Authors:** Muhammad Asif, Hugo F.M.C. Martiniano, Ana Rita Marques, João Xavier Santos, Joana Vilela, Celia Rasga, Guiomar Oliveira, Francisco M. Couto, Astrid M. Vicente

## Abstract

The complex genetic architecture of Autism Spectrum Disorder (ASD) and its heterogeneous phenotype make molecular diagnosis and patient prognosis challenging tasks. To establish more precise genotype-phenotype correlations in ASD, we developed a novel machine learning integrative approach, which seeks to delineate associations between patients’ clinical profiles and disrupted biological processes inferred from their Copy Number Variants (CNVs) that span brain genes. Clustering analysis of relevant clinical measures from 2446 ASD cases in the Autism Genome Project identified two distinct phenotypic subgroups. Patients in these clusters differed significantly in ADOS-defined severity, adaptive behaviour profiles, intellectual ability and verbal status, the latter contributing the most for cluster stability and cohesion. Functional enrichment analysis of brain genes disrupted by CNVs in these ASD cases identified 15 statistically significant biological processes, including cell adhesion, neural development, cognition and polyubiquitination, in line with previous ASD findings. A Naive Bayes classifier, generated to predict the ASD phenotypic clusters from disrupted biological processes, achieved predictions with a high Precision (0.82) but low recall (0.39), for a subset of patients with higher biological Information Content scores. This study shows that milder and more severe clinical presentations can have distinct underlying biological mechanisms. It further highlights how machine learning approaches can reduce clinical heterogeneity using multidimensional clinical measures, and establish genotype-phenotype correlations in ASD. However, predictions are strongly dependent on patient’s information content. Findings are therefore a first step towards the translation of genetic information into clinically useful applications, but emphasize the need for larger datasets with very complete clinical and biological information.

## Introduction

Autism Spectrum Disorder (ASD) is a neurodevelopmental disorder that manifests with persistent deficits in social communication and interaction, and unusual or repetitive behaviour and/or restricted interests [1]. ASD presents a highly heterogeneous clinical phenotype and frequently co-occurs with other comorbidities, such as Intellectual Disability (ID), epilepsy and Attention Deficit Hyperactivity Disorder (ADHD) [2–6]. Heritability estimates indicate a strong genetic influence in ASD aetiology [7–9], however reliable genetic markers for the disease are unavailable. ASD is diagnosed through neurodevelopmental assessment, which can be challenging especially in the case of very young children. Improving early diagnosis and prognosis using biological markers with a robust predictive power would provide an advantage to young patients, who benefit the most from an early start of specific intervention [10].

Copy Number Variant (CNV) screening is nowadays widely used for etiological diagnosis, with causative genetic alterations identified in approximately 25% of ASD cases [11]. A large number of rare genetic variants have been implicated in ASD, and the wide genetic heterogeneity that characterizes this disorder likely contributes to phenotypic variability in ASD patients [12]. Integrative pathway and network analysis of large scale ASD genomic studies have advanced significantly the identification of disrupted biological processes [13–17]; however, our understanding of the biological meaning of the large number of putative pathogenic variants, their phenotypic manifestations, and the reliable interpretation of many genetic findings for clinical application is still lagging.

To improve our ability to infer clinical meaning from rare CNVs in ASD, for eventual application as biological markers, we developed a machine learning-based approach involving the integration of gene functional annotations and clinical phenotypes. Our approach was developed in four steps, namely: 1) definition of clinically distinct subgroups in ASD cases; 2) discovery of functionally enriched biological processes defined by rare CNVs disrupting brain-expressed genes in the same ASD cases; 3) assessment of the contribution of disrupted biological processes for classification of ASD phenotypes; 4) design and predictive effectiveness characterization of a machine learning classifier for clinical outcome in ASD patients.

## Methods

Figure 1 shows the graphical representation of the overall methodology, described in detail below.

**Figure 1:**
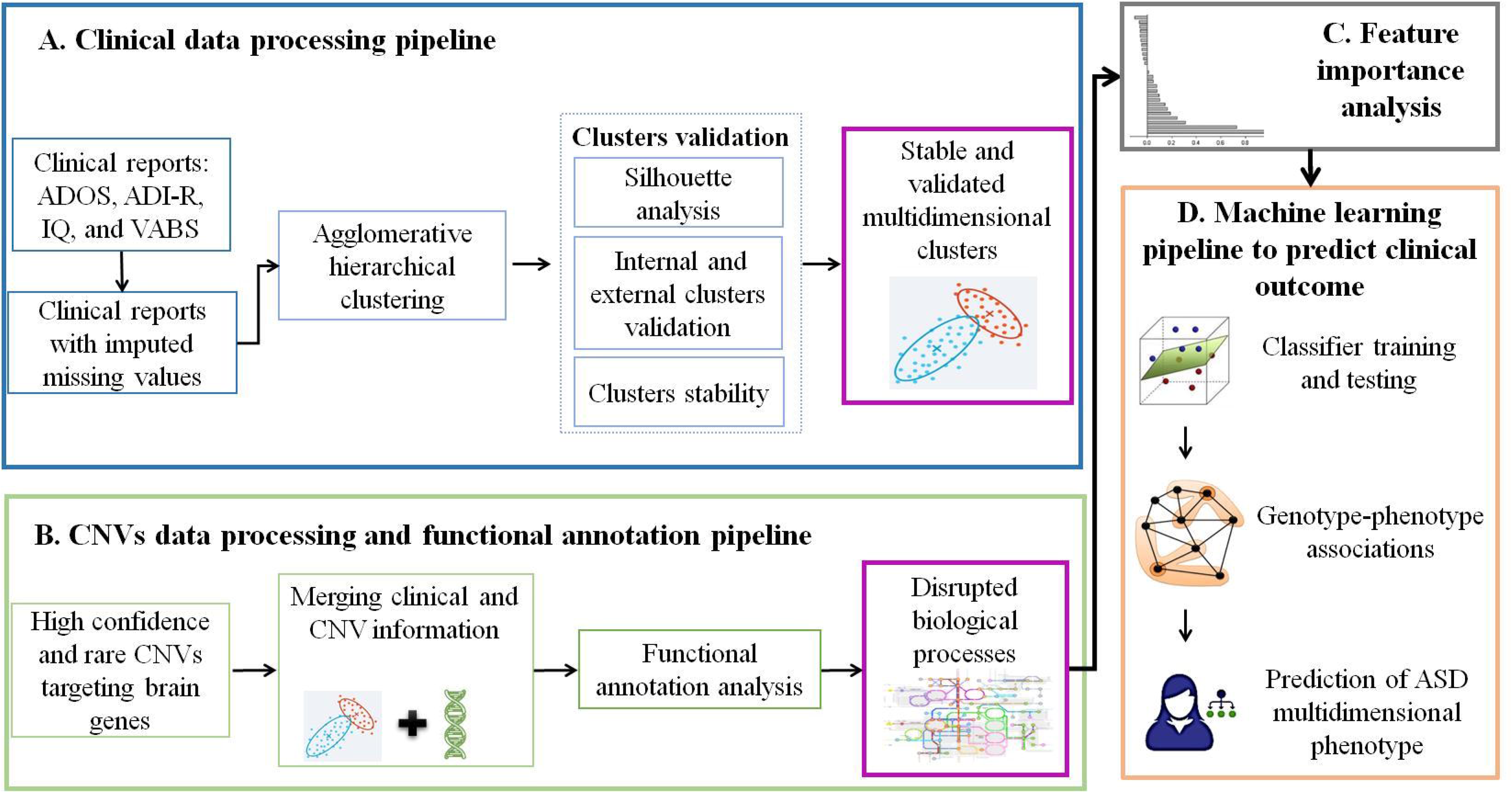
Integrative systems medicine approach to identify complex genotype-phenotype associations. Clinical and genetic data from the Autism Genome Project (AGP) was used in this study (A) Clinical data analysis processing: clinical data comprises reports of ASD diagnosis and neurodevelopmental assessment instruments. Agglomerative Hierarchical Clustering (AHC) was used to identify clinically similar subgroups of individuals in stable, validated clusters, defined by multiple clinical measures. (B) CNV data processing: rare high confidence CNVs previously identified by the AGP, targeting brain-expressed genes, were retained for analysis. CNV data was merged with clinical data from clustered ASD subjects for a final list of CNVs targeting brain genes. (C) Functional annotation analysis: Biological processes defined by brain-expressed genes targeted by CNVs were obtained using g:Profiler. (D) Classifier design: A Naive Bayes machine learning classifier was trained and tested on patient’s data, to predict the phenotypic clustering of patients from biological processed disrupted by rare CNVs targeting brain-expressed genes.

### Participants

The ASD dataset used in this study was obtained from the Autism Genome Project (AGP) [18] database, and comprises CNV data and clinical information from 2446 ASD patients. The AGP was an international collaborative effort from over 50 different institutions to identify risk genes for ASD. The group of individuals with phenotypic information from clustering and rare CNV data, used in final analysis included 1213 males (83.4%) and 144 females (10.6%).

### ASD diagnosis, clinical assessment instruments and clinical features

Individuals meeting criteria defined by the Diagnostic and Statistical Manual of Mental Disorders IV (DSM–IV) [19] and the thresholds for Autism or ASD from the Autism Diagnostic Interview-Revised (ADI-R) [20] and the Autism Diagnostic Observation Schedule (ADOS) were classified as ASD cases [21]. The AGP defined a phenotypic classification system based on the combined ADI-R and ADOS diagnosis, categorizing subjects into Strict (meeting thresholds for Autism by the ADI-R and ADOS), Broad (meeting thresholds for Autism from one instrument and ASD from the other) and Spectrum (meeting thresholds for Autism from at least one instrument or ASD from both). Individuals with an ASD diagnosis from only one instrument and no information from the other, or not meeting thresholds for Autism or ASD from one of the instruments, regardless from the classification from the other, were not included in the study. Clinical measures used in this study were retrieved from the AGP database, including the ADIR verbal status, ADOS severity score, Vineland Adaptive Behaviour Scales (VABS) [22] subscales and an Intelligence Quotient (IQ).

The ADI-R verbal status is a dichotomized measure indicating the verbal status of the individual at evaluation. The ADOS severity metric ranges from 1 to 10 and is calculated from ADOS modules 1 to 3 raw scores [23]. As there is no algorithm available to calculate ADOS severity score for ADOS module 4 reports, which is applied only to adolescents and adults, subjects with the ADOS module 4 (N= 149) were dropped from further processing. The severity score distribution is collapsed into three categories, namely Autism (severity scores ranging from 6 to 10), ASD (severity scores ranging from 4 to 5) and Non-Spectrum (severity scores from 1 to 3), which reflect the mapping of the severity metric onto raw ADOS scores. The ADOS Non-spectrum category includes individuals with a mild phenotype, and in this study 125 individuals with a Non-spectrum ADOS severity score fell within the Spectrum phenotypic class from the AGP, meaning they met thresholds for autism from the ADI-R, and were thus included.

The VABS is used to assess adaptive functioning of individuals and consists of three subscales, namely, socialization, communication and daily living skills scores, and also computes a composite score. Subjects with VABS scores ≤70 were classified in a dysfunctional adaptive behavior category, for all subscales. IQ scores of ASD cases were also retrieved from the AGP database, and categorized with the following thresholds: IQ>70 normal, 50<IQ<70 mild intellectual disability, IQ<50 severe intellectual disability.

Clinical reports from the ASD patients were examined for missing values, and clinical features with more than 70% information were retained for the analysis. To minimise missing value imputation bias, individuals with missing values above this threshold for more than two clinical features were also excluded. Completeness of each clinical feature is reported in Table S1 (Additional file 1). Missing values were imputed using the missForest [24] R package that implements the Random Forest [25] algorithm, a decision tree-based supervised machine learning method. Imputation error was assessed using the normalised global Proportion of Falsely Classification (PFC), and the missing values imputation error was 0.12.

### Clustering analysis of ASD clinical data

To focus on core domains of ASD symptoms, verbal skills, disease severity, adaptive behavior and intellectual levels, which strongly condition prognosis, were selected for further analysis. Verbal status was obtained from the ADI-R, ASD severity scored from the ADOS, adaptive functioning from the VABS, using its three subdomains, and a performance IQ category from the IQ assessment contributed by participating sites to the AGP database. Other IQ domains had too many missing values to be used. The Agglomerative Hierarchical Clustering (AHC) [26] method was used to identify independent phenotypic subgroups from the selected clinical features. Correlations between clinical features were assessed using the Pearson method, and features with a correlation value of > 0.75 were considered correlated. The Gower [27] metric was used to calculate the distance matrix from the patient’s clinical data. To normalise the effect of highly correlated variables on clustering, the weight for correlated variables (VABS subscales of socialisation, communication, and daily living skills) was reduced to half during distance matrix calculation. To identify phenotypic subgroups, the AHC method using Ward2 [28] criteria was applied to the distance matrix.

To assess the contributions of each clinical feature in defining the clusters, we excluded one feature at a time, re-performed the clustering and observed the changes in Silhouette values of both clusters. For this purpose, we selected Silhouette value as an evaluation metric because it was also used to define outliers in clinical data. A decrease in the Silhouette value of a cluster after removing one feature indicates its importance in defining this cluster and vice versa.

### Goodness of clustering assessment

A Silhouette method [29] was employed to estimate the goodness of the clustering results. The Silhouette value for each individual shows how well the individual is clustered, and ranges from −1 to 1, with individuals scoring below 0 considered as wrongly clustered. In addition, the Silhouette value for each cluster was derived, and clusters with Silhouette value of > 0.25 were considered as true clusters. Bootstrapping with 1000 iterations was used to measure the stability of clusters, where a boot mean value above 0.85 corresponds to stable clusters. All clustering analysis was performed in R environment, using Cluster [30] and FPC packages.

### Functional enrichment analysis

Genotyping and CNV calling methods for the AGP ASD subjects (N=2446) were previously described [18]. CNVs called by any two algorithms (high confidence CNVs) and above 30kb in size were retained for further analysis. To screen for rare CNVs (<1% in control population) CNV frequencies in control populations were estimated using the genotypes from the studies by Sheikh et al. [31] (N = 1320) and Cooper et al. [32] (N = 8329), identified using the same genotyping platform [18]. Control genotypes were obtained from the Database of Genomic Variants (DGV) [33].

To focus CNV selection on variants spanning brain-expressed genes, avoiding *a priori* hypotheses from ASD candidate gene assumptions, an extensive list comprising 15585 brain-expressed genes was obtained from Parikshak et al. [34]. The brain-expressed gene list was prepared from brain RNA-seq data, collected at thirteen different developmental stages, including genes expressing during early brain developmental phase. The full criteria and parameters used to define the brain-expressed gene list were previously described [34].

The g:Profiler [35] tool was employed to identify biological processes enriched for brain-expressed genes spanned by rare CNVs in ASD individuals. g:Profiler implements a hypergeometric test to estimate the statistical significance of enriched biological processes, followed by multiple corrections for the tested hypotheses using the Benjamini-Hochberg procedure. g:Profiler uses Gene Ontology (GO) data to find the biological annotations for input genes.

The GO tool contains a Directed Acyclic Graph (DAG) structure with a clear hierarchical parent-to-child relationship between GO terms. Because of this DAG structure, functional enrichment analysis can result in redundant GO terms, which may lead to high correlations between GO terms. To minimise the correlations between GO terms, the Revigo [36] tool was employed to redundant GO results. Revigo uses the methods of semantic similarity to measure similarities between GO terms. The SimRel [37] method was used to calculate similarities between GO terms, and terms with a similarity score of > 0.7 were grouped.

### Feature importance assessment

The mean decrease in accuracy of the Random Forest algorithm was used to compute the importance score of each disrupted biological process for categorizing ASD subjects into defined phenotypic clusters. A stratified ten-fold cross-validation quantifies the importance of all features. The importance score of all disrupted biological processes was recorded at each fold. A final importance score for each biological process was calculated by averaging their importance score values across all the ten folds. Random Forest was implemented using randomdForst R package [38].

### Classifier learning and cross-validation

A Naive Bayes [39] machine learning method was employed to predict the ASD phenotypic group, defined by the clustering analysis, from biological processes disrupted by rare CNVs. This method employs the Bayes theorem of probability for training and testing of the model, and the algorithm was implemented using the klaR R package with default parameters. Precision, recall, specificity and F-score were used as evaluation measures. To train and test the Naive Bayes, a stratified five-fold cross-validation approach was used, in which data was first split into five equal subsets with equal class probabilities; a Naive Bayes model was trained on any four subsets, and the remaining subset was used as the test set. This process was repeated five times and each time a different subset was used as test set. For each repetition, the model performance was estimated and mean values for precision, recall, specificity and F-score were reported. The Naive Bayes classifier was trained on patient’s data by using the “more severe” cluster as the positive class and the “less severe” cluster as the negative class.

The Information Content (IC) from each individual represents the level of specificity of biological processes disruption, and was derived by summing the IC values of all the biological processes disrupted in each individual. IC is a numerical value that describes the specificity of a GO term using its position in the GO DAG structure.

## Results

### Identification of ASD clusters defined by clinical phenotype

A total of 1817 ASD subjects from the AGP were retained for analysis after assessment of missing values in clinical features. Agglomerative hierarchical clustering analysis of clinical observations from these patients initially identified two phenotypically independent clusters. To minimise the phenotypic complexity and define the most stable and cohesive clusters, weakly clustered individuals with a Silhouette value less than 0.300 (representing a balance between number of individuals lost and goodness of clustering) were excluded from the clustering analysis. After removal of weakly or wrongly clustered individuals, cluster 1 contained 903 ASD cases, while cluster 2 comprised 494 patients (Table 1). Elimination of the loosely clustered individuals resulted in more stable and cohesive clusters, with high values for clusters stability and reduced average distance between the two individuals in a cluster (Table 1).

**Table 1:** Clustering validation, after removal of weakly clustered individuals.

| Clusters validation measures | Cluster 1 | Cluster 2 |
| --- | --- | --- |
| Clusters size (N) | 903 | 494 |
| Average distance between two patients | 0.235 | 0.231 |
| Silhouette value | 0.567 | 0.579 |
| Average Silhouette of both clusters | 0.571 |  |
| Cluster stability | 0.998 | 0.996 |

Overall, the cluster validation through the Silhouette method and bootstrapping showed that both clusters were true and consistent.

### Clinical interpretations of the clusters

All clinical measures differed significantly between the two clusters, as shown in Table 2. Cluster 1(Additional file 1: black circles in Figure S1) includes a higher number of individuals, who generally exhibited a milder clinical phenotype, while Cluster 2 (Additional file 1: red triangles in Figure S1) included a higher percentage of subjects with severe dysfunction. All individuals in Cluster 1 were verbal according to the ADI-R, while Cluster 2 included only non-verbal cases. The mean age of ADI-R assessment was 7.7 years, an age when verbal status is generally well established. Furthermore, the mean age of individuals in Cluster 1 (mean age 8.02) and Cluster 2 (mean age 7.01) did not significantly differ.

**Table 2:** Clusters 1 and 2 statistics for each clinical measure.

| Clinical measure | Clinically defined categories | Cluster 1 N (%) | Cluster 2 N (%) | *p-value* |
| --- | --- | --- | --- | --- |
| ADIR verbal status | ADI-R-non verbal | 0 (0) | 494 (100) | <0.00001<sup>a</sup> |
|  | ADI-R-verbal | 903 (100) | 0 (0) |  |
| ADOS severity score | ADOS severity score Autism (score 6-10) | 714 (79.07) | 392 (79.35) | <0.00001<sup>b</sup> |
|  | ADOS severity score ASD (score 4-5) | 64 (7.09) | 102 (20.65) |  |
|  | ADOS severity score Non-spectrum (score 1-3) | 125 (13.84) | 0 (0) |  |
| VABS communication | Dysfunctional VABS communication (score ≤ 70) | 307 (34) | 493 (99.8) | <0.00001<sup>a</sup> |
|  | Normal VABS communication (score > 70) | 596 (66) | 1 (0.2) |  |
| VABS daily living skills | Dysfunctional VABS daily living skills (score ≤ 70) | 478 (52.94) | 484 (97.98) | <0.00001<sup>b</sup> |
|  | Normal VABS daily living skills (score > 70) | 425 (47.07) | 10 (2.02) |  |
| VABS socialization | Dysfunctional VABS socialization (score ≤ 70) | 497 (55.04) | 490 (99.19) | <0.00001<sup>a</sup> |
|  | Normal VABS socialization (score > 70) | 406 (44.96) | 4 (0.81) |  |
| Performance IQ Scale | Severe disability (score <50) | 2 (0.22) | 218 (44.13) | <0.00001<sup>b</sup> |
|  | Moderate disability (score ≥ 50 and ≤70) | 31 (3.43) | 125 (25.3) |  |
|  | Normal ability (score > 70) | 870 (96.35) | 151 (30.57) |  |
| Gender | Male | 830 (91.92) | 417 (84.41) | 0.000015<sup>b</sup> |
|  | Female | 73 (8.08) | 77 (15.59) |  |

<sup>a</sup>Fisher Exact Test, <sup>b</sup>Chi-Square test

For all VABS sub-domains, roughly half of the subjects in Cluster 1 were in the normal range; conversely, over 97% of individuals belonging to Cluster 2 showed dysfunctional adaptive behaviour. Consistent with the other clinical measures, over 96% of cases from Cluster 1, but less than one third in Cluster 2, scored at the normal level in performance IQ, while a much higher percentage of ASD cases from Cluster 2 than from Cluster 1 presented with a performance IQ in the range of severe intellectual disability.

Regarding the ADOS severity score, approximately 14% of the individuals in Cluster 1 were assigned to the milder category of the ADOS severity score (“Non-spectrum” for ADOS, but scoring positive for “Autism” in the ADI-R, and therefore classified in the AGP “Spectrum” phenotypic class, see methods). Conversely, none of the individuals in Cluster 2 scored in this category. On the other hand, a significantly higher percentage of cases in Cluster 2 (20.65%) than individuals in Cluster 1 (7.09%) scored in the intermediate ASD severity category. It is noteworthy that both clusters show a similarly high percentage of individuals scoring in the “Autism” ADOS severity category. This is not surprising since this broad category (scores ranging from 6 to 10) comprises all subjects classified in the Strict AGP phenotype class but also a large proportion of individuals in the AGP Broad phenotype class. The “Autism” ADOS severity score therefore targets a subset of the study population that can be quite heterogeneous in phenotypic presentation. Corroborating this, we found that the “Autism” category of the ADOS severity score is not significantly associated with the clusters (*χ*2= 0.15, *p* = 0.901, df = 2), even though overall there is a significant association of the overall ADOS severity scores (Table 2).

Both clusters were strongly dominated by the male gender, partly because of the high percentage of males in the dataset after the elimination of weakly or wrongly clustered individuals. However, the percentage of males was higher in cluster 1, representing the milder phenotype, consistent with general observations that male to female ratios are higher in datasets that comprise more high-function ASD individuals.

Analysis of the contribution of each clinical feature in defining clusters showed that the main contributor was the ADIR verbal status variable (Additional file 1: Table S2). The VABS subscales had a strong effect on Cluster 1 but a modest role in defining Cluster 2. Performance IQ also contributed more to Cluster 1 whereas for Cluster 2 it has the least effect. The ADOS severity score did not have a major role in defining either cluster, as indicated by the similar high percentage of subjects scoring within the range of “Autism” in the ADOS severity scale in both clusters. Similarly, gender was not an important contributor to the definition of either cluster.

### Disrupted biological processes from brain-expressed genes targeted by rare CNVs

CNVs (N=129754) identified in 2446 subjects with ASD were filtered to select rare, high confidence CNVs, over 30 Kb in size and that contained complete or partial brain-expressed gene sequences. The selected high confidence, rare CNVs (N=12683) disrupted 4025 brain-expressed genes in 2414 subjects with ASD (86.8% males and 13.2% females).

Phenotypic cluster and rare CNV data was complete for 1357 individuals with ASD, and available for integration. Functional enrichment analysis of rare CNVs targeting brain-expressed genes (N=2738) in 1357 patients identified 17 statistically significant biological processes (Additional file 1: Table S3). g:Profiler did not recognize 187 genes from the input list.

The redundancy of GO terms in functional enrichment analysis, caused by overlapping annotations in ancestors and descendent terms in the DAG structure of GO, was reduced by grouping the terms that had a semantic similarity score higher than 0.7 (Additional file 1: Table S3). The Revigo tool used to reduce redundancy did not recognise one biological process (*Plasma membrane bounded cell projection organization*). After redundancy reduction, 16 biological processes remained (Table 3), with the *Calcium-dependent cell-cell adhesion via plasma membrane cell adhesion molecules* biological process merged with *Homophilic cell adhesion via plasma membrane adhesion molecules* (similarity score = 0.76).

**Table 3:** Statistically significant enriched biological processes for CNVs spanning brain-expressed genes (N=2738). FDR: False Discovery Rate

| Biological processes | Enriched genes (N) | FDR *p*-value |
| --- | --- | --- |
| Homophilic cell adhesion via plasma membrane adhesion molecules | 53 | 6.30E-09 |
| Cell-cell adhesion via plasma-membrane adhesion molecules | 66 | 1.70E-07 |
| Cellular component organization or biogenesis | 944 | 5.70E-05 |
| Cellular component organization | 915 | 7.00E-05 |
| Cellular component biogenesis | 475 | 0.00066 |
| Cellular component assembly | 434 | 0.00177 |
| Nervous system development | 363 | 0.00215 |
| Organelle organization | 562 | 0.00475 |
| Protein polyubiquitination | 64 | 0.00592 |
| Cell projection organization | 231 | 0.00836 |
| Cellular localization | 418 | 0.0091 |
| Single-organism behavior | 83 | 0.0196 |
| Regulation of cellular component organization | 364 | 0.0257 |
| Plasma membrane bounded cell projection organization | 223 | 0.0282 |
| Cognition | 56 | 0.0364 |
| Single-organism organelle organization | 263 | 0.044 |

The most significant biological process identified in this dataset was *Homophilic cell adhesion via plasma membrane adhesion molecules*, which includes 53 brain-expressed genes disrupted by the selected CNVs. The ten most significant biological processes were related to cell adhesion and cellular organization, and also included nervous system development and protein poliubiquitination (Table 3). Moreover, two significant biological processes were related to behavior and cognition.

### Biological process importance for prediction of ASD clinical phenotype

The enriched biological processes and phenotypic cluster information for ASD cases were combined in a matrix to assess the predictive value of the biological processes for categorization in one of the two phenotypic clusters, broadly characterized by a milder and a more severe phenotypic presentation. The 57 individuals containing both rare CNV and cluster information that did not present any enriched biological process were excluded, so further analysis comprised 1300 ASD patients.

Table 4 shows the ranking in importance of disrupted biological processes for categorization of subjects into ASD phenotypic clusters, computed using the Random Forest importance score function.

**Table 4:** Importance of each biological process from Random Forest in classifying ASD subjects into defined phenotypic clusters.

| Random Forest rank | Biological process | Mean Decrease in Accuracy |
| --- | --- | --- |
| 1 | Regulation of cellular component organization | 0.052 |
| 2 | Cell projection organization | 0.025 |
| 3 | Cellular component assembly | 0.025 |
| 4 | Single organism behaviour | 0.020 |
| 5 | Organelle organization | 0.018 |
| 6 | Single organism organelle organization | 0.017 |
| 7 | Cellular component biogenesis | 0.014 |
| 8 | Cognition | 0.013 |
| 9 | Nervous system development | 0.010 |
| 10 | Cellular localization | 0.009 |
| 11 | Cellular component organization | 0.006 |
| 12 | Protein polyubiquitination | 0.005 |
| 13 | Homophilic cell adhesion via plasma membrane adhesion molecules | 0.005 |
| 14 | Cell adhesion via plasma membrane adhesion molecules | 0.005 |
| 15 | Cellular component organization or biogenesis | 0.003 |

The importance of each biological process was calculated using the mean decrease in accuracy, computed by permuting each biological process. The feature importance analysis using Random Forest, which was trained and tested using stratified 10-fold cross-validation over the integrated dataset, revealed positive values for all features, indicating that all of the biological processes are positively contributing for classification. The most important biological process for the classification was *Regulation of cellular component organization*, with a mean decrease in accuracy of 0.052. The most significantly enriched biological process in the overall ASD dataset, *Homophilic cell adhesion via plasma membrane adhesion molecules* was ranked at position 14, indicating it is not a top contributor to phenotypic categorization of ASD subjects into the phenotypic clusters, in this population.

### Predicting clinical phenotype from the biological processes disrupted by rare CNVs in ASD patients

The Naive Bayes supervised machine learning method was trained and tested using phenotypic clustering information and the 15 biological processes inferred from rare CNVs targeting brain-expressed genes in ASD patients. The classifier was trained with the assumption that ASD subjects with a more dysfunctional clinical phenotype, subgrouped in Cluster 2, would present a different pattern of disrupted biological processes from the individuals with a milder expression of ASD phenotype in Cluster 1.

The Naive Bayes classifier trained on data from 1300 patients did not perform well in predicting the more dysfunctional clinical phenotype from disrupted biological processes (Table 5), with scores indicating a low accuracy of the predictive model.

**Table 5:** Naive Bayes performance in predicting the severe phenotype of ASD

| Data used for classification | N | Precision | Recall | Specificity | F-score |
| --- | --- | --- | --- | --- | --- |
| All ASD cases | 1300 | 0.221 | 0.379 | 0.655 | 0.279 |
| ASD cases from 1st quantile with highest IC | **325** | **0.816** | **0.389** | **0.699** | **0.526** |
| ASD cases from 1st and 2nd quantiles of IC | 649 | 0.23 | 0.384 | 0.65 | 0.284 |
| ASD cases from first three quantiles of IC | 974 | 0.29 | 0.389 | 0.672 | 0.329 |

To further dissect the information available, the biological process Information Content (IC) for each individual was calculated by summing the IC values for all the biological processes disrupted in that individual. ASD subjects in the first IC quantile (N = 325) had highest IC scores, while ASD cases belonging to fourth quantile (N = 326) contained lowest IC scores. The performance of the Naive Bayes classifier improved when only ASD subjects with higher IC were selected for analysis. Analysis of the group of individuals with highest IC (first quantile) resulted in a higher predictability of ASD clinical outcome (Table 5). The classifier trained and tested on individuals from the first two (1^st^ and 2^nd^) and first three (1^st^, 2^nd^, and 3^rd^) quantiles also performed better than the classifier designed using the whole dataset of clusters and biological processes (Table 5).

The Naive Bayes classifier was thus able to make reasonably good predictions of ASD severity, but only for a subset of ASD individuals with higher IC. This indicates that improved GO information, as well as larger datasets with more GO information available, are needed to usefully integrate clinical and biological data.

## Discussion

The discovery of diagnostic and prognostic biomarkers for ASD has the potential to improve the reliability of diagnosis at earlier stages of development, as well as the phenotypic categorization for prognosis, eventually informing personalized intervention that is particularly beneficial for very young children. However, in spite of the enormous volume of genetic information generated by genomic approaches in the past decade, the clinical diagnosis of ASD patients is still solely based on neurodevelopmental assessment. The results of many genomic tests, including CNV arrays and clinical exomes, still leave about 80% of the cases without any explanation regarding the biological pathways underlying their disease and their personal clinical presentation.

In this study, we developed a novel integrative approach to predict ASD phenotypes from biological processes defined by genetic alterations. Overall, our approach sought to exploit multidimensional clinical measures to define subgroups of ASD patients presenting similar clinical profiles, and then to identify the biological processes disrupted by CNVs that might predict these more homogeneous clinical patterns. For the sake of eventual clinical utility, we chose clinical measures with well-established relevance and frequently used in clinical settings, but established no other restrictions. Further, we did not set any *a priori* hypothesis for gene selection, besides being expressed in the brain.

The clustering of clinical data from ASD cases resulted in two subgroups that were clearly distinguishable in terms of severity of phenotype, defined by multiple clinically relevant measures including verbal status, ASD severity, adaptive function and cognitive ability. The identification of only two clusters for the clinical phenotype, with an important proportion of individuals in the AGP dataset that could not be adequately clustered was expected, as it reflects the high clinical heterogeneity of ASD. The identification of these subgroups was in line with previous results by Veatch et al. [40], who also identified two clusters differing in severity using two independent population samples, including the Autism Genetic Resource Exchange (AGRE) and also the AGP dataset. While clinical variables were not fully coincidental between the two studies, we confirmed that the verbal status, ADOS-based severity, VABS-based communication, socialization and daily living skills, as well as gender, were all significantly different between clusters. We noted an unequal contribution of each clinical measure to definition of each cluster, with verbal status the main contributor and the ADOS severity score a low contributor for both clusters, while Performance IQ was mainly important for Cluster 1.

In our study, the larger Cluster 1 was characterized by a generally milder phenotype, with all individuals being verbal, a large proportion in the normal IQ range and significantly higher numbers of subjects scoring better in adaptive behavior subscales. Cluster 1 also showed a higher male to female ratio, as expected given the general observation that higher functioning ASD subgroups have a larger proportion of males. The smaller Cluster 2 included only non-verbal subjects, and had a higher percentage of subjects with a more dysfunctional phenotype in terms of adaptive behavior, as well as lower IQ scores. Because cognitive ability is such an important variable for prognosis, we included performance IQ as a clinical variable, in spite of the limitations related to the heterogeneity of IQ measurement tools used for patient assessment by AGP contributing sites. For the AGP dataset, an effort was previously made to rationalize the tests used, and cognitive level was established using a categorical classification provided by AGP sites in three categories, namely severe intellectual disability, mild intellectual disability and normal IQ, for verbal, performance and full scale IQ scores. Limitations were also introduced by the proportions of missing data; given the adopted control of the validity of imputation procedures, only performance IQ met the criteria for reliable imputation, so only this measure was used.

Because our main goal was to improve the power for phenotypic subgroup prediction by genetically defined biological processes, we focused on obtaining compact and stable clusters by using strict criteria for cluster stability to assess the goodness of clustering, at the expense of population sample dimension. As expected, the weakly clustered individuals tended to have more divergent scores across clinical measures (data not shown), and therefore were more difficult to cluster with high confidence. It is intriguing that a higher proportion of females than males was removed, suggesting that this divergence of scores is more frequent in girls. This observation generally supports recent debates on the lower adequateness of assessment criteria to the female autism phenotype [41].

To test the hypothesis that phenotypic subgroups have specific underlying pathological mechanisms, we first sought to identify the biological processes enriched in the gene sets disrupted by rare CNVs detected in the AGP dataset. The functional enrichment analysis conducted in this study was independent of any prior assumptions or weighting criteria of genes relative to ASD risk. To make functional enrichment analysis hypothesis-free and to let the data speak, we screened for CNVs disrupting any brain-expressed genes. The objective was to obtain a complete picture of the convergence of rare CNVs, targeting any brain-expressed genes, into biological processes relevant for brain function.

The biological processes identified in the functional enrichment analysis showed an overlap with putative core biological mechanisms of ASD defined by previous studies. For example, 363 brain genes spanned by rare CNVs were enriched for neurodevelopment biological process and 56 genes were associated with cognition process. Enrichment of nervous system development and cognition processes in ASD has been previously reported by studies using different approaches, including transcriptome analysis and co-expression networks [15] and is supported by the function of genes most consistently implicated in ASD, like *PTEN, RELN, SYNGAP1, ANK2, SCN2A* and *SHANK3* [42]. Noh et al. analysis of *de novo* CNVs spanning ASD genes also implicated cognitive processes, and showed a convergence in cellular component organization or biogenesis, cellular component assembly, and organelle organization biological processes [16]. Other studies implicated cell adhesion processes in ASD as important components of synapse formation and function (46, 47). Dysregulation of polyubiquitination was also in line with previous studies reporting an excess of variants in genes involved in ubiquitination processes, which regulate neurogenesis, neuronal migration and synapse formation, and are thus essential for brain development [43–46].

This biological heterogeneity parallels the extensive phenotypic heterogeneity that characterizes ASD. For this reason, we sought to identify the biological processes underlying the more homogeneous phenotypic subgroups defined by the clusters. The Random Forest algorithm was used to assess the importance of each enriched biological process in discriminating the two ASD phenotype subgroups. Feature importance analysis showed that all the biological process contributed positively to the classification of ASD severity. However, the feature importance ranking was different from the significance ranking of enriched biological processes. Despite their relevance for ASD, the top three statistically significant biological processes identified by functional enrichment analysis were least important for the classification of subjects into the phenotypic milder and more dysfunctional subgroups. These findings support the concept that the integration of datasets with multidisciplinary information, including genomic and clinical data, is necessary to discover the biological mechanisms that lead to specific clusters of symptoms.

The Naive Bayes classifier was able to make useful predictions of ASD phenotypic subgroups from disrupted biological processes, but only for a subset of individuals for whom annotations had higher information content for the biological processes defined by the CNVs. Currently, GO contains more than 40,000 biological concepts, which are rapidly evolving with the increasing knowledge of biological phenomena and with our ability to structure this knowledge. Therefore, it is expected that the performance of the proposed classifier will improve with the progress in GO annotations.

Given the high clinical heterogeneity of ASD, clustering of individuals according to a multidimensional phenotype will result in subgroups with more homogeneous clinical patterns and for whom the causes of this disease are more likely to have the same underlying biological mechanism. The clustering of individuals according to multidimensional clinical symptoms *per se* is likely to have implication for prognosis and outcomes, as concurrent symptoms may have a synergistic effect on disease progression, and may thus also help guide clinical practice and intervention. However, thus far this perspective has been insufficiently explored, and not enough datasets are yet available with detailed clinical information that can be merged for large scale analysis. The alterations in diagnostic criteria over time and the changes in versions of instruments like the ADI-R and the ADOS create important challenges for data merging across population samples, which are needed so that sufficient statistical power is achieved for definite conclusions. This study is clear in this limitation, as the number of subjects with important missing data in multiple clinical features was high in the AGP dataset, reducing analytical power, and thus only two stable clusters could be defined. The next research steps will necessarily have to involve overcoming limited clinical information and merging challenges between available datasets, like AGRE and the Simons Foundation Autism Research Initiative (SFARI), so that models established for biological predictions can be useful in clinical settings. On the other hand, while genomic information gets easier and cheaper to collect, improvements are also necessary regarding GO annotations; a large number of subjects with phenotypic subgroup data did not have sufficient GO information content to be useful for classifier predictions.

## Conclusion

Overall, the present approach is proof of concept that genotype-phenotype correlations can be established in ASD, and that biological processes can predict multidimensional clinical phenotypes. Importantly, it highlights the usefulness of machine learning approaches that take advantage of multidimensional measures for the construction of more homogeneous clinical profiles. It further stresses the need to overcome the limitations of analyzing individual gene variants in favor of considering biological processes disrupted by an heterogeneous set of gene variants. The results stress two major requisites for translation of genomic information into useful clinical applications: that study datasets include detailed and complete clinical information, and that databases containing biological process information are rigorously and extensively curated. Identification of biological processes for specific clinical subgroups will be important to discover physiological targets for pharmacological therapy that can be efficient for subgroups of patients. This strategy can equally become very useful in clinical settings, for predicting outcomes and planning interventions for subgroups of patients whose specific patterns of clinical presentation are defined by the genes disrupted by specific genetic variants.

## Acknowledgements

Work was supported by UID/MULTI/04046/2013 centre grant from Portuguese Fundação para a Ciência e Tecnologia (FCT), Portugal (to BioISI), UID/CEC/00408/2019 (LASIGE) and PTDC/CCI-BIO/28685/2017 (DeST: Deep Semantic Tagger Project) to FMC, and MA was recipient of a fellowship from BioSys PhD programme (Ref: SFRH/BD/ 52485/2014) from FCT (Portugal). Patients and parents were genotyped in the context of the Autism Genome Project (AGP), funded by NIMH, HRB, MRC, Autism Speaks, Hilibrand Foundation, Genome Canada, OGI, and CIHR. We acknowledge the families who participated in these projects.

