## Additional file for "Identification of biological mechanisms underlying a multidimensional ASD phenotype using machine learning"

Table S1: Completeness of each clinical feature after removing individuals with uncertain diagnosis and with ADOS module 4 reports. Total number of individuals = 1892

| Clinical feature | Completeness (%) |
| --- | --- |
| Gender | 100 |
| AGP phenotype class (strict, broader, spectrum, and not included) | 100 |
| ADIR verbal status | 99.10 |
| VABS socialization | 97.04 |
| VABS communication | 96.30 |
| VABS daily living skills | 96.09 |
| VABS composite score | 95.61 |
| Seizures | 89.90 |
| ADOS severity score | 79.44 |
| Performance IQ Scale | 73.84 |
| Verbal IQ scale | 63.79 |
| Full scale IQ | 53.75 |

Table S2: Contribution of each clinical feature in defining clusters.

| Clinical feature | Cluster 1 | | | Cluster 2 | | |
| --- | --- | --- | --- | --- | --- | --- |
|  | Silhouette | Average distance between two individuals | Stability | Silhouette | Average distance between two individuals | Stability |
| ADIR verbal status | 0.364 | 0.3 | 0.785 | 0.461 | 0.264 | 0.675 |
| Performance IQ | 0.451 | 0.271 | 0.999 | 0.752 | 0.138 | 0.998 |
| Gender | 0.584 | 0.254 | 0.998 | 0.639 | 0.224 | 0.996 |
| VABS daily living skills | 0.612 | 0.216 | 0.986 | 0.553 | 0.246 | 0.974 |
| VABS communication | 0.612 | 0.214 | 0.998 | 0.525 | 0.254 | 0.997 |
| VABS socialization | 0.63 | 0.209 | 0.999 | 0.546 | 0.253 | 0.998 |
| ADOS severity score | 0.636 | 0.209 | 0.639 | 0.643 | 0.209 | 0.639 |

Table S3: Functionally enriched biological processes for rare CNVs disrupting 2738 brain-expressed genes in 1357 patients and their semantic similarity scores. FDR: False Discovery Rate, NA: Not Available

| Biological processes | Enriched genes (N) | FDR *p*-value | Semantic similarity score |
| --- | --- | --- | --- |
| Protein polyubiquitination | 64 | 0.00592 | 0 |
| Homophilic cell adhesion via plasma membrane adhesion molecules | 53 | 6.3E-09 | 0 |
| *Calcium-dependent cell-cell adhesion via plasma membrane cell adhesion molecules* | *12* | *0.037* | *0.765* |
| Single-organism behavior | 83 | 0.0196 | 0 |
| Cognition | 56 | 0.0364 | 0 |
| Cellular localization | 418 | 0.0091 | 0 |
| Cellular component organization or biogenesis | 944 | 5.7E-05 | 0 |
| Cellular component organization | 915 | 7E-05 | 0.004 |
| Nervous system development | 363 | 0.00215 | 0.167 |
| Cell projection organization | 231 | 0.00836 | 0.359 |
| Single-organism organelle organization | 263 | 0.044 | 0.388 |
| Regulation of cellular component organization | 364 | 0.0257 | 0.42 |
| Cellular component assembly | 434 | 0.00177 | 0.443 |
| Cellular component biogenesis | 475 | 0.00066 | 0.458 |
| Organelle organization | 562 | 0.00475 | 0.479 |
| Cell-cell adhesion via plasma-membrane adhesion molecules | 66 | 1.7E-07 | 0.617 |
| Plasma membrane bounded cell projection organization | 223 | 0.0282 | NA |

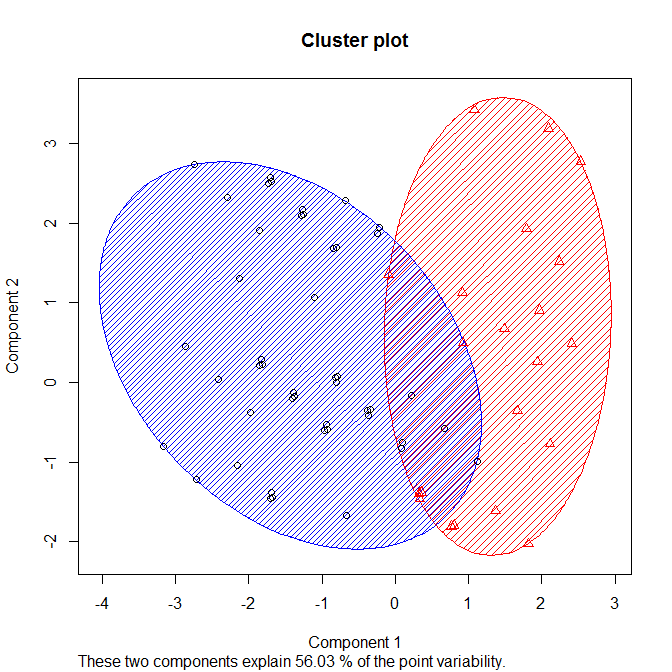

Figure S1: Subgrouping of ASD patients. Black circles enclosed by blue ellipse represent cluster 1 individuals and red triangles enclosed by red ellipse are cluster 2 individuals
